# Orally administered niclosamide-based organic/inorganic hybrid suppresses SARS-CoV-2 infection

**DOI:** 10.1101/2022.07.19.500639

**Authors:** Geun-woo Jin, Goeun Choi, N. Sanoj Rejinold, Huiyan Piao, Young Bae Ryu, Hyung-Jun Kwon, In Chul Lee, Jin-Ho Choy

**Author notes:** **Correspondence** (J.-H.C.). (H.P.), (N.S.R.).

## Abstract

The COVID-19 pandemic is a serious global health threat mainly due to the surging cases along with new variants of COVID-19. Though global vaccinations have indeed some effects on the virus spread, its longevity is still unknown. Therefore an orally administrable anti-viral agent against SARS-CoV-2 would be of substantial benefit in controlling the COVID-19 pandemic. Herein, we repurposed niclosamide (NIC), an FDA approved anthelmintic drug in to MgO, which was further coated with hydroxyl propyl methyl cellulose (HPMC) to get the de-sired product called NIC-MgO-HPMC, which has improved anti-SARS-CoV-2 replication in the Syrian hamster model. The inhibitory effect of NIC-MgO-HPMC on SARS-CoV-2 replication leads to the prevention of inflammation as well as lung injury. These data strongly support that repurposed NIC-MgO-HPMC could be highly beneficial for controlling the ongoing pandemic thereby achieving an endemic phase.

## 1. Introduction

The COVID-19 disease has been labelled as pandemic in March 11, 2020. [1-3]. SARS-CoV-2 is a highly transmissible virus and a surging cases worldwide has threatens human health and public safety overwhelming the healthcare systems. So far, it affected an estimate of ∼ 561 M individuals all around the world with ∼ 6.37 M deaths as of Jul 16, 2022. Vaccines were expected to make this pandemic controllable, but SARS-CoV-2 virus remains a part of our lives due to the emerging new variants such as the delta [4], omicron, B.1.1.529 [5], deltacron [6] and the latest BA.2.75 (Centaurus), etc., making the current situation as emergency to come up with new anti-SARS-CoV-2 therapeutics as a better option to treat COVID-19 patients, than vaccination alone.

As Merck & Co. and Pfizer Inc. have started clinical trials for their experimental anti-SARS-CoV-2 pills such as Lagevrio (Molnupiravir) [7-10] and Paxlovid [11, 12], the race has begun toward an efficient and convenient COVID-19 therapeutics. However, the lack of long-term safety data concerning the new drugs could be an obstacle. For example, Lagevrio works as faulty RNA building block when the virus utilizes polymerase enzyme to copy its genome. In addition, this kind of drugs were found to be mutagenic even to the mammalian cells as well [13]. The Phase 3 analysis of Lagevrio includes criteria such as refraining from sperm donation and agreement to abstaining from intercourse otherwise use contraceptives. In addition, female groups had to be neither pregnant nor breastfeeding. Additionally, women of childbearing age must have had negative pregnancy test within 24 h before the first dose of medication. Another concern is the high price of the drug, ∼ 700 USD per course as per the recent Merck contract with the US government [14]. The biggest problem with existing antiviral drugs is that they cannot cope with newly emerging mutated variants. Direct acting antivirals, such as Paxlovid or Lagevrio, have an inherent limitation that their anti-viral activity could be lowered when a structural change in the target viral enzyme occurs.

In addition, on May 31st 2022, the G7 academies have jointly stated the need for a new broad spectrum anti-viral agent and “be prepared” for the next pandemic in advance. Present FDA approved drugs have been suffering from drug resistance for the new variants and there is an urgent need for broad spectrum anti-viral agent which can be used for wide range of viruses such as paramyxoviruses, bunyaviruses, togaviruses, filoviruses (including Ebola viruses and Marburg virus), picornaviruses (including enteroviruses and other cold-causing viruses), and flaviviruses (including the viruses that cause yellow fever, dengue and Zika) which are potential to cause future pandemics.

In this context, niclosamide (NIC) is an excellent antiviral drug candidate since NIC has proven its broad-spectrum antiviral activity through previous studies however, the reason of the broad spectrum antiviral activity has not been clarified yet. In 2019, Gassen et. al., found that NIC induces autophagy, enabling a mechanistic explanation of the broad spectrum antiviral activity of NIC. In 2021, it was found that SARS-CoV-2 virus was also inhibited in proliferation due to autophagy induced by NIC through a study by the same institute [15, 16].

However, the major trouble is the low aqueous solubility, thereby poor bioavailability (∼10%) of NIC that could limit its potential effect on the SARS-CoV-2 virus [17] since the systemic absorption of NIC would be troubled by the oral administration. For instance, when an existing digestible tablet was orally administered, only a negligible plasma concentration was observed for anti-SARS-CoV-2 replication [8]. One of the major approaches for enhancing the drug solubility, is through chemical modification by which NIC could have better therapeutic effects [18]. Since such chemically modified NIC derivatives are new ones, they have to undergo a thorough preclinical and clinical studies to assure their safe usage. The other approach is a drug repurposing technology [17] by which one can enhance bioavailability of NIC through either physical and/or chemical modification with pharmaceutically inert excipients. This is believed to fuel the drug development process faster since the toxicological profiles are well established [19].

We have learned from long-term research that inorganic materials can improve the bioavailability of drugs [20, 21]. It was hypothesized that a large specific surface area of inorganic compound, such as hydrotalcite (HT) could contribute to the improved bioavailability of hydrophobic drugs [22]. Based on this idea, we invented the NIC oral formulation using MgO, which is an FDA approved antacid having a rock salt structure. In addition, MgO itself could be used as an adjuvant medicine for COVID-19 therapy as evidenced by several reported studies [23]. The NIC-MgO oral formulation was finally coated with pharmaceutically inert, FDA approved excipient, hydroxy propyl methyl cellulose (HPMC) [24-28], to make the final product as NIC-MgO-HPMC which is expected to have better pharmacokinetic (PK) effects to sustain the NIC release for improved therapeutic effects against SARS-CoV-2 pre-clinical and clinical studies . Previously, we could also understand that repurposed NIC formulation with montmorillonite (MMT) [29], hydrotalcite [22], porous silicas and geopolymer [30] enhanced the PK performance of NIC. In addition, all the main components, such as the MgO, NIC and HPMC have been approved by the FDA, making the present NIC-drug formulation, a safe one for anti-COVID therapy.

Further, the anti-viral efficacy study using SARS-CoV2-infected hamster model involved a medically applicable dosage range. Several previous trials have tested anti-viral efficacy with clinically inapplicable drug dosage range. For example, the antiviral activity of Lagevrio and Paxlovid was orally administered to mouse models (Balb/c nude) with 250 mg/kg [31,32] and 300 mg/kg, respectively. Human equivalent doses (HED) calculated based on allometric scale are 2.4g and 2.9g for Lagevrio and Paxlovid respectively. Herein, we showed that newly repurposed NIC oral formulation, NIC-MgO-HPMC has an ability to inhibit SARS-CoV-2 infection in hamster model with a dosage below 100 mg/kg. In reality, the developed oral NIC formulation on administration, inhibits SARS-CoV-2 infection in hamster model at dose of 40 and 80mg/kg. These obtained information aided a foundation for investigational new drug (IND) application data for NIC-MgO-HPMC, which is now in clinical trial Phase 2.

## 2. Materials and Methods

### 2.1. Materials

NIC was purchased from DERIVADOS QUIMICOS. Ethanol was acquired from Daejung Chemicals. MgO was purchased from Sigma-Aldrich (Korea). HPMC (6 mPa·s) was gifted by Wonpoong Pharm Co., Ltd, South Korea.

### 2.2. Preparation of NIC-MgO and NIC-MgO-HPMC

The NIC-MgO and NIC-MgO-HPMC were prepared by conventional solid-state intercalation reaction. The NIC-MgO was synthesized by mixing NIC and MgO with weight ratio of NIC : MgO = 4 : 2.8, and grinding with 70% ethanol. The NIC-MgO-HPMC was synthesized by mixing NIC, MgO and HPMC with weight ratio of NIC : MgO : HPMC = 4 : 2.8 : 1, and grinding with 70% ethanol.

### 2.3. Preparation of CTAB (Cetrimonium bromide) stabilized MgO and NIC-MgO NPs

For CTAB stabilized NPs, initially CTAB was dissolved in distilled water under heating at 45°C for 2 min, then 1:1 ratio (CTAB: Samples ∼ MgO, NIC-MgO) were used for the synthesis. Briefly, 5 ml of CTAB (1 mg/mL) was added with 5 mg of powder samples and then mixed gently at 300 rpm for1 15 min, until the samples were well dispersed in the CTAB solution. The samples were then analyzed for the DLS analysis after probe sonicating for 20 sec and then syringe filtered using 450 nm PVDF membrane.

### 2.4. NIC content analysis

The drug content in the samples were analysed by treating with 99.9% EtOH, followed by 30 min probe-sonication thereafter NIC content was extracted fully. Final steps involved, sample filtration using a 450 nm sized PVDF filter, and thereafter absorption peaks were detected at 333 nm using UV spectrophotometer (V-630, Japan).

### 2.5. *In-vitro* drug release study

The drug release test was done according to the reported work using a DST-810 dissolution apparatus (Lab fine, Korea), at fixed temperature of 37 °C with a stirring frequency of 50 rpm. The NIC release was done by in two different solutions with pH 1.2 and 6.8, both of them containing 2% Tween 80 in it, to simulate the human’s gastric and intestinal fluids (USP, 2002a, pp. 2011-2012). Samples containing identical NIC content (18 mg) were suspended in the dissolution chamber. Thereafter, the aliquots were taken at pre-scheduled time points. The drug release was quantified by measuring characteristic NIC absorption at 333 nm using UV spectroscopy.

### 2.6. Characterizations

For FT-IR analysis, a Jasco FT-IR-6100 spectrometer (JASCO, Tokyo, Japan) was used with the standard KBr disk method in transmission mode (spectral range 4000–400 cm^-1^, resolution 1 cm^-1^, 40 scans per spectrum). For FT-IR, 100 mg of KBr pellet was mixed with 12 mg samples, ground well, pelletized using hydraulic press machine (20 MPa for 1 min). Particle size and surface charge measurements were done using Otsuka Electronics DLS/Zeta EL-SZ-2000 (Japan) using disposable square cuvettes in distilled water. The powder X-ray diffraction (XRD) measurements were conducted using a Bruker D2 Phase diffractometer (Bruker, Karlsruhe, Germany) equipped with Cu Kα radiation (λ = 1.5418 Å). All the data were recorded at the tube voltage and a current of 30 kV and 10 mA, respectively. The UV–Visible absorption for the NIC, NIC-MgO and NIC-MgO-HPMC were measured in EtOH (99.9%) at 333 nm by a Jasco UV/Vis spectrometer (V-630, Easton, MA, USA).

### 2.7. PK study in hamsters

This experiment was done on golden Syrian hamsters (male, 11-weeks aged, 129.55–160.93 g). The animal experiments were sanctioned by KNOTUS Institutional Animal Care and Use Committee (KNOTUS IACUC 21-KE-771). Prior to oral administration of the NIC oral formulation, the powder was dispersed in water. The blood samples were centrifuged for 2 min at 13,000 rpm in order to separate the plasma and frozen immediately for HPLC-mass spectrometry (HPLC-MS) analysis.

### 2.8. NIC Quantification hamster’s plasma

Plasma calibration curve was made by different NIC concentrations of 0, 5, 10, 50, 100, 500, 1000, and 2000 ng/mL. Samples were prepared by adding 100 µL of the internal standard, topiramate, to 20 μL aliquot of hamster’s plasma sample. Then, after 5 sec vortexing, centrifugation was done (13,000 g for 5 min at 4 °C). The 20 µL of supernatant was treated with 180 µL of 50 % methanol. Thereafter 150 µL of the samples were analysed by Acquity I-Class UPLC system (Waters) with mass spectrometer (Waters Xevo TQ-S). The LC analytical column was a Thermo HGypersil Gold (2.1 × 50 mm, 1.9 microns), which was maintained at 50°C throughout the analyses and the mobile phase B was made of 5mM ammonium acetate water solution, and methanol. The flow rate was 0.4 mL/min and the gradient was 50% A initially, then later at 1.5 min, it was increased to 70 % B. From 1.5 min to 2.5 min, the mobile phase was maintained at 70 % B and at 4.0 min was switched immediately back to 50 % B.

### 2.9. Anti-viral efficacy study

Hamsters (11 week-old) were divided into groups for 20, 40, 80mg/kg of NIC-MgO-HPMC and NIC treatment (n = 6 per group). Groups were then treated with the oral formulations dissolved in water at 6 h following infection (post-infection). 12 h dosing regimen was retained till the completion of the study in day 4 post-infection. A third group consisted of vehicle control animals that received the same dosing schedule and volume as NIC-MgO-HPMC treated groups. All groups were infected intranasally with 2 × 10^5^ PFU/mL of SARS-CoV-2. Animals were sacrificed on day 4 post-infection and lung tissues were collected at necropsy for pathological and virological analyses.

### 2.10. Viral load

RNA was extracted from homogenized lung tissues using the RNeasy Mini kit (Qiagen) according to the manufactures protocol. For detection of viral RNA, it was used in a one-step real-time RT-PCR against the N gene using primer (Forward: 5’-TAA TCA GAC AAG GAA CTG ATT A-3’ // Reverse: 5’-CGA AGG TGT GAC TTC CAT G-3’) and CFX96 Touch Real-Time PCR Detection System (Bio-Rad).

### 2.11. Histopathology and immunohistochemistry

Histopathology and immunohistochemistry were performed on hamster lung tis-sues. Tissues were fixed in 10 % Neutral Buffered Formalin with two changes, for a minimum of 7 days. Tissues were placed in cassettes and processed with a Sakura VIP-6 Tissue Tek, on a 12-h automated schedule, using a graded series of ethanol, xylene, and paraffin embedded tissues were sectioned at 5 µm and dried over-night at 42°C prior to staining. Lung injury score was calculated as below

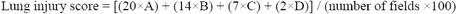

(where, A: inflammatory cell infiltration in alveolar space, B: Inflammatory cell infiltration in interstitial space, C: hyaline membrane, D: Haemorrhage)

### 2.12. Gross lung pathology

Hamsters were assigned a clinical score based on signs of disease as determined by gross pathology of lungs

### 2.13. Toxicology study

The toxicology study of NIC-MgO-HPMC was conducted by administrating the formulation for 7 days (bid, 12h interval). The animals were monitored for 14 days (7 days of dosing period+7 days of recovery period). Sprague-Dawley (NTacSam:SD) rats (6-week-old, male: 158.8∼194.1g, female: 132.9∼164.7g) and Beagle dogs (5-7 months old, male: 6.78∼8.62kg, female: 5.60∼7.58kg) were used to assess the toxicity of NIC-MgO-HPMC formulation. The toxicology studies were sanctioned by Dt & CRO Institutional Animal Care and Use Committee (SD rat: 210281, Beagle dog: 210215). On day 15, animals were sacrificed to observe abnormalities in organs.

### 2.14. Statistical information

Statistical analysis was performed in R. The difference in viral load and gross lung lesion between study arms was assessed by One-way ANOVA followed by a Kruskal-Wallis test and a pairwise Wilcoxon rank sum test to correct for multiple comparisons. The lung injury score from histopathological study between study arms was evaluated by Kruskal-Wallis rank sum test and a pairwise Wilcoxon rank sum test to correct for multiple comparisons.

## 3. Results

The PXRD patterns for various samples such as NIC, MgO, NIC-MgO, HPMC, and NIC-MgO-HPMC were analyzed to confirm the possible intercalation of NIC within MgO. The PXRD pattern of MgO showed specific peaks corresponding (111), (200), (220) reflection planes, indicating the cubic crystalline structure for the MgO, confirming the characteristic periclase phase for MgO (Database file no. 45-0946). The PXRD results clearly indicated that NIC was incorporated very well in the MgO and further coating with HPMC resulted no obvious changes in the PXRD pattern (Fig.1a). Further, energy dispersive elemental analysis revealed the presence of all the essential components within the composite particles (Fig. 1b).

**Fig. 1.**
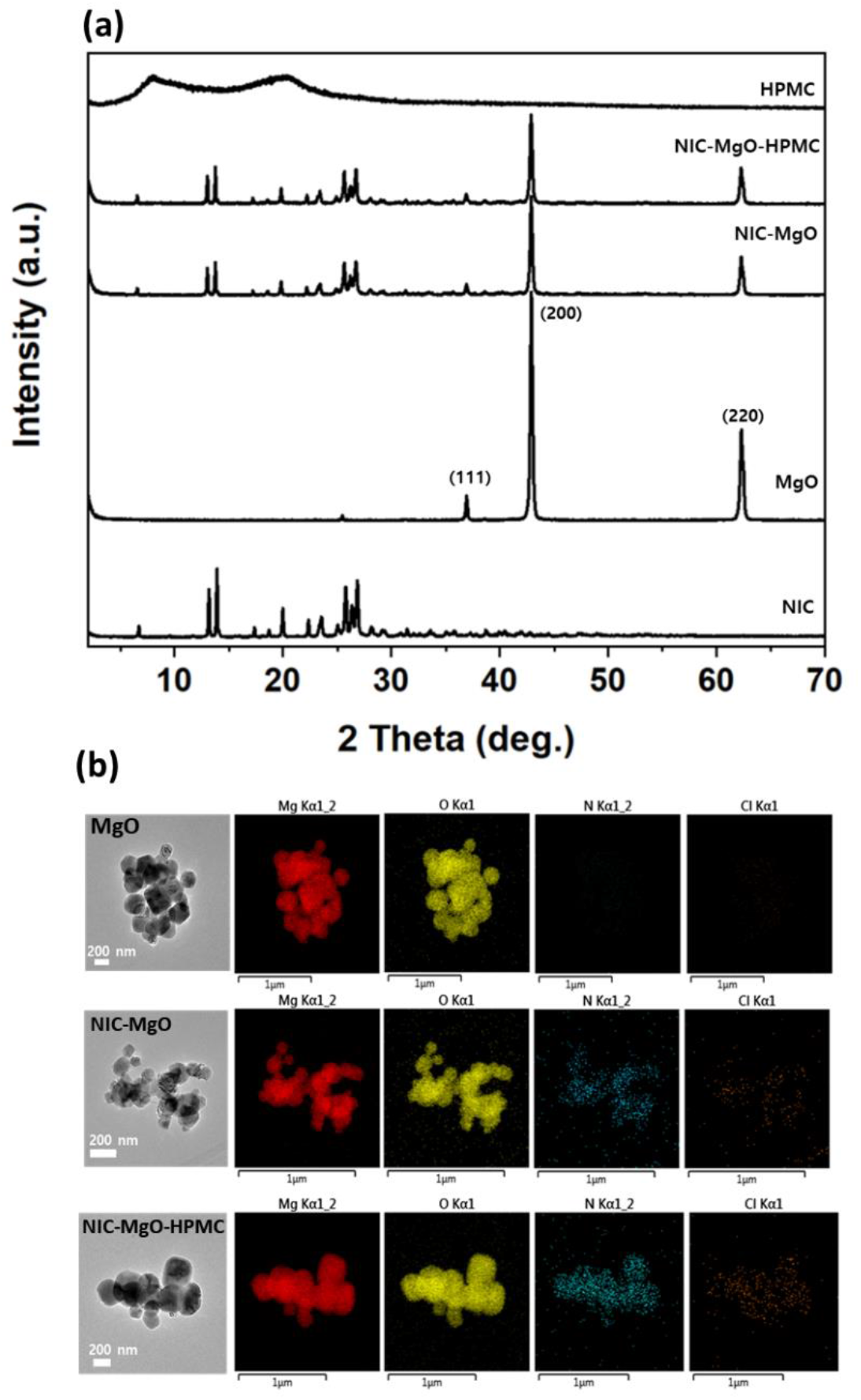
(a) PXRD patterns of pristine NIC, MgO, NIC-MgO, NIC-MgO-HPMC and HPMC. (b) TEM images and EDS mapping analyses of MgO, NIC-MgO, NIC-MgO-HPMC.

The FE-SEM analyses were done on samples such as NIC, MgO, NIC-MgO, NIC-MgO-HPMC in order to distinguish their morphological transformation as such and their composite forms (Fig. 2). As previously studied, NIC showed rod like morphology [22] whereas the recrystallized NIC exhibited almost a featureless morphology, but highly disordered one. On the other hand, MgO NPs showed spherical morphology having a particle size of 200 nm. These results were in well agreement with the TEM analysis as shown in Fig.1b.

**Fig. 2.**
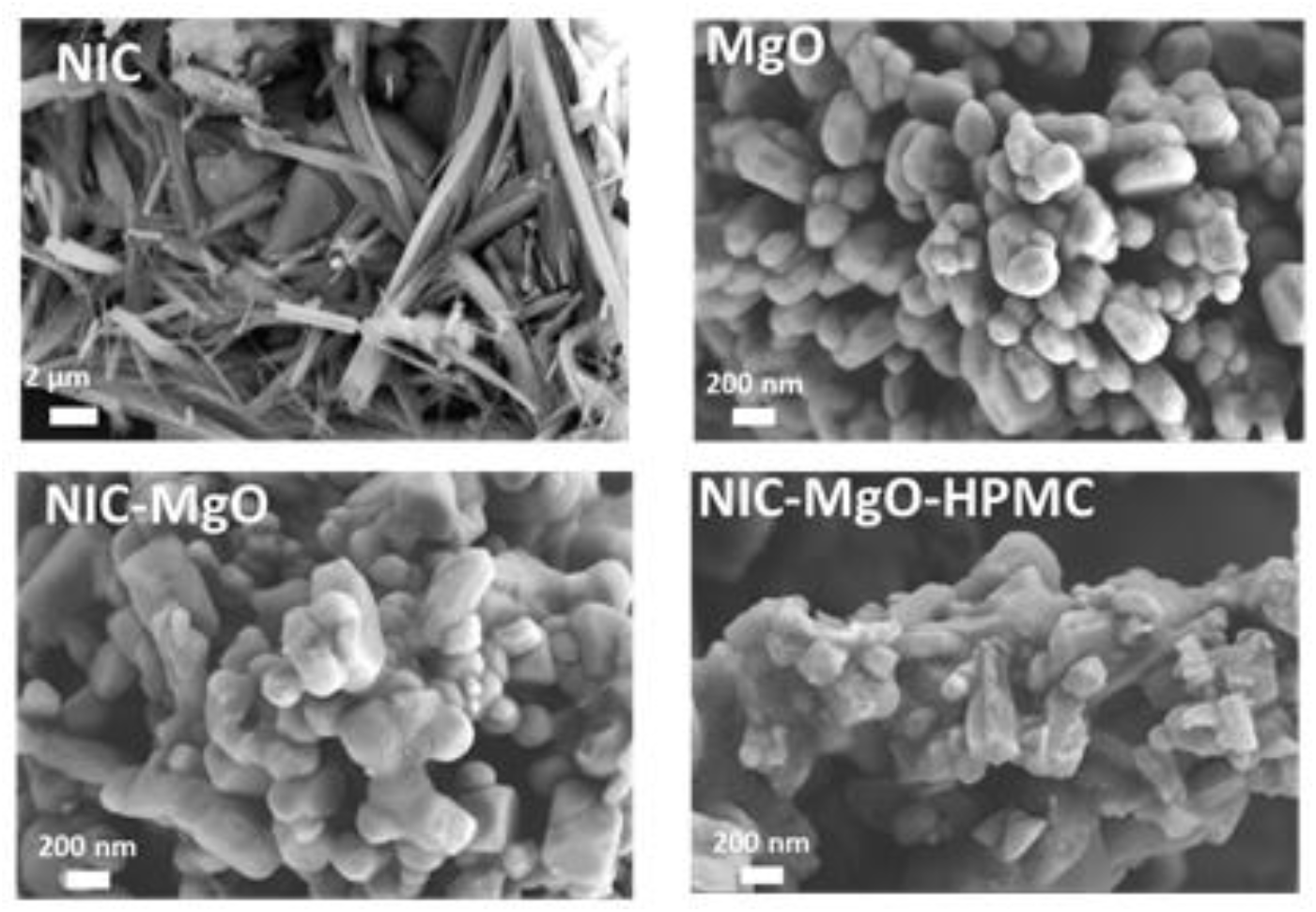
**(a)** FE-SEM images of NIC, MgO, NIC-MgO and NIC-MgO-HPMC.

Figure 3 shows the Fourier transform infrared (FT-IR) spectra of intact NIC, NIC-MgO and and NIC-MgO-HPMC. The intact NIC showed major characteristic peaks at 3578 cm^-1^, 3490 cm^-1^, 1680 cm^-1^, 1510 cm^-1^, and 540 cm^-1^, those which could be as-signed as -OH, -NH, -C=O, -NO_2_, and C–Cl groups, respectively. For MgO, broad bands around at 3400cm^-1^ was observed, which can be ascribed to (O–H) group vibrations. These characteristic peaks were vanished in both NIC-MgO and NIC-MgO-HPMC samples indicating rather there must be a strong chemical interaction within their nanohybrids.

**Fig. 3.**
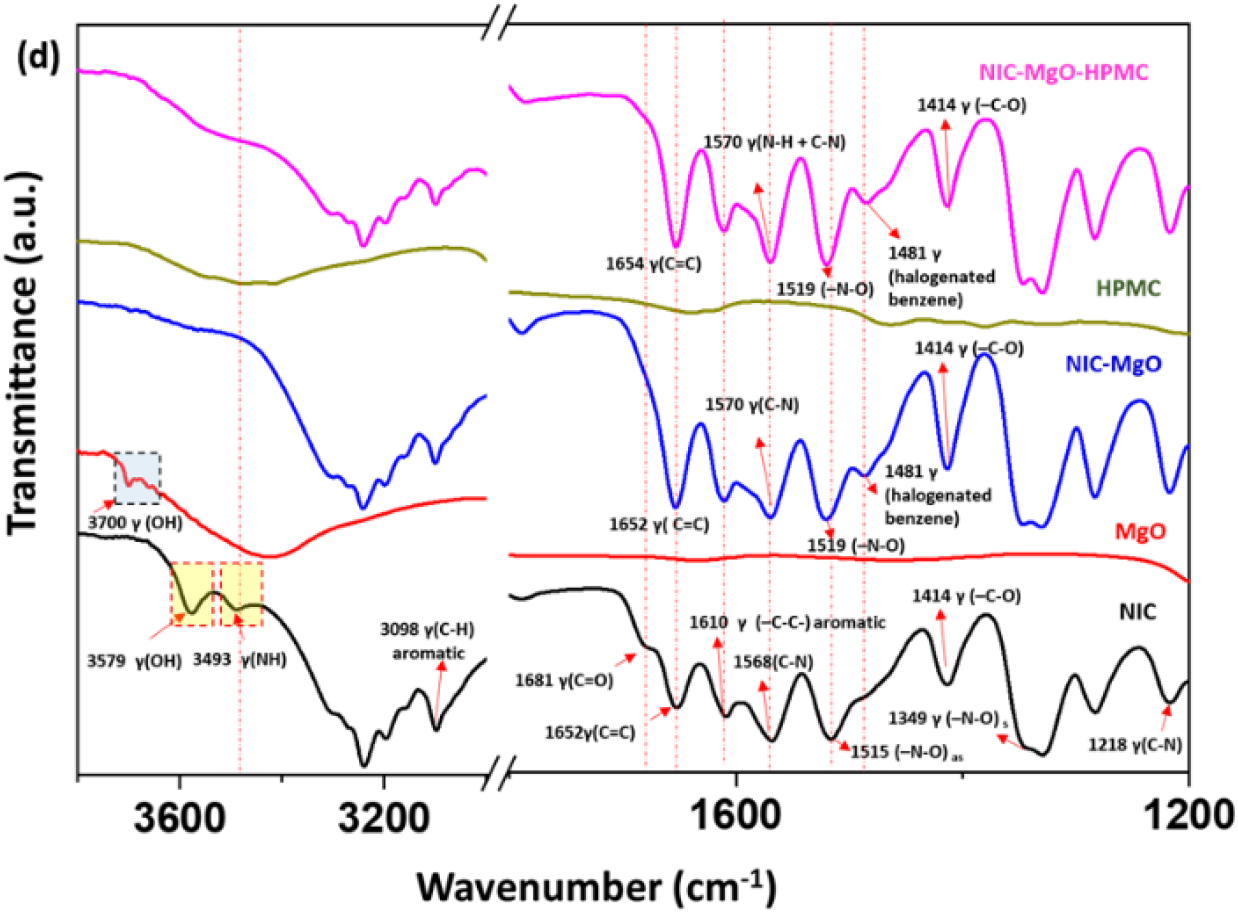
FT-IR spectra for NIC, NIC-MgO and NIC-MgO-HPMC.

The DLS and zeta analyses were carried out in distilled water to understand the particle size distribution (Fig. 4). The MgO particles were not well dispersed in water, therefore we coated initially with CTAB (Cetrimonium bromide) in 1:1 ration with MgO and NIC-MgO in order to get stable particles in the distilled water. The cationic CTAB has a chain length of 1.5 to 2 nm [33] and could easily be coated on the negatively charged MgO particles. Accordingly, the CTAB-MgO NPs had size of ∼ 222.3 ± 8.3 nm almost similar to the TEM results, whereas the NIC-MgO NPs showed a size of 245.9 ± 9. 9 nm. Since the chain length of CTAB is very small, its coated samples are expected show no significant differences, meaning that the both MgO and NIC-MgO NPs must be in the range ∼ 220 nm. The HPMC coated NIC-MgO, however, showed the particle size of 214.4 ± 3.1 nm, clearly due to the stabilization via HPMC coating [34]. In addition, all the samples have shown PDI values < 0.4 (Table 1), whereas, the HPMC coated one had better aqueous dispersity among others.

**Table 1.** DLS, Zeta and PDI measurements for various samples such as CTAB stabilized MgO NPs, NIC-MgO and HPMC coated NIC-MgO in distilled water. The HPMC coated samples show better stability in terms of PDI.

| Samples | Size (nm) | Zeta (mV) | PDI (a.u.) |
| --- | --- | --- | --- |
| **MgO (intact)** | 330.7±7.8 | -22.24 ± 2.2 | 0.341 ± 0.027 |
| **MgO-CTAB** | 222.3 ± 8.3 | 42.2 ± 3.6 | 0.360 ± 0.03 |
| **NIC-MgO-CTAB** | 245.9 ± 9.9 | 42.1 ± 3.4 | 0.345 ± 0.025 |
| **NIC-MgO-HPMC** | 214.4 ± 3.1 | 21.4 ± 5.8 | 0.396 ± 0.013 |

**Fig. 4.**
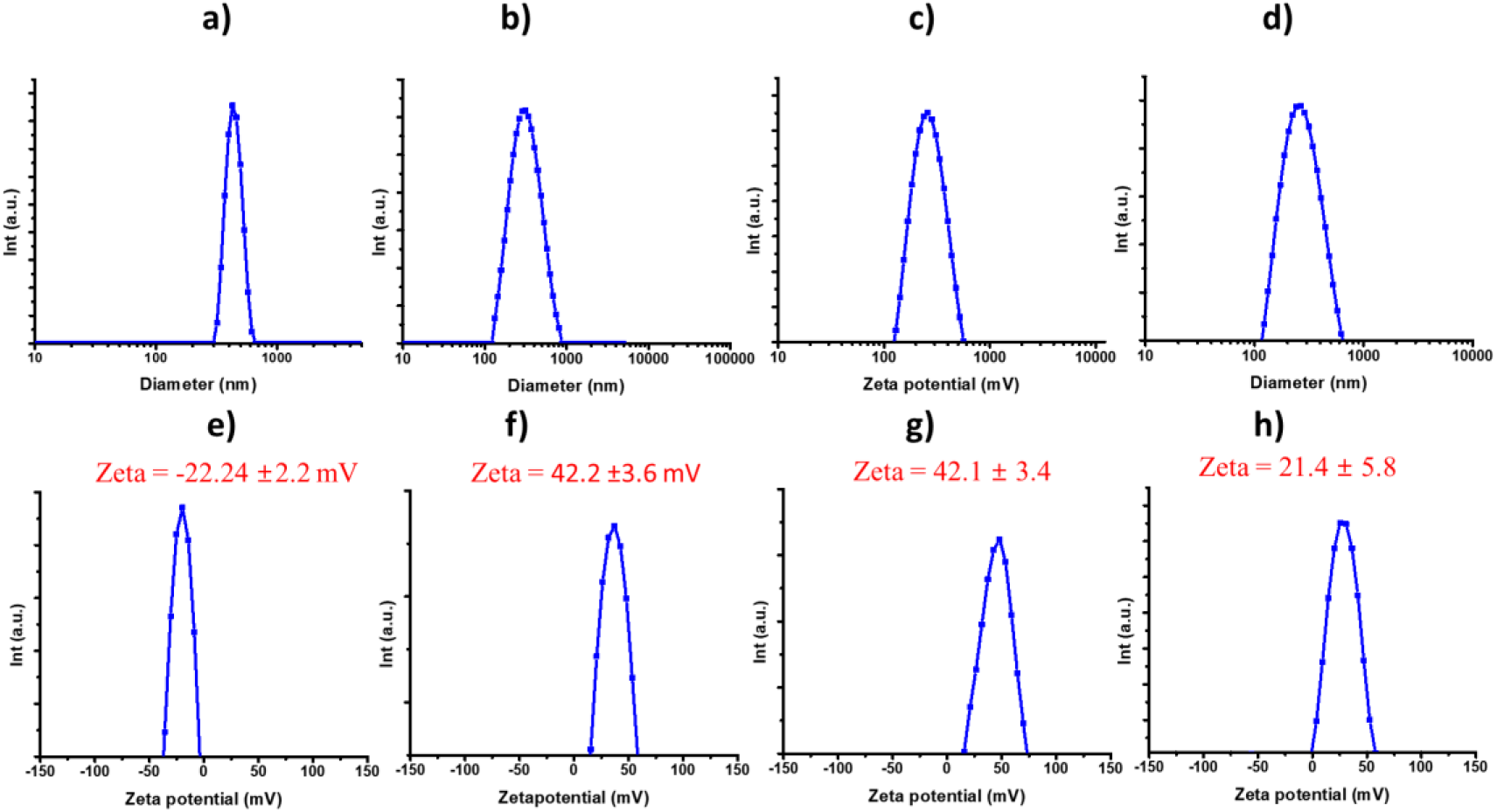
DLS analysis for a) Intact MgO, b) CTAB-MgO, c) CTAB-NIC-MgO, d) NIC-MgO-HPMC, where e) to h) are their corresponding zeta measurements in distilled water.

In general, there are two types of conformations found in NIC, which are of phenyl-phenyl orientation via π–π antiparallel and π–π parallel-displaced stacking. In addition, it was reported that the NIC cluster size could be associated with either parallel or antiparallel π–π stacking, which was equally observed from the equilibrium system. A metastable parallel conformation would be dominant if there is an increased NIC monomers in the clusters, followed by the T-shaped conformation, which resembles to those found in proteins. Such micro clusters can be stabilized by strong hydrophobic π–π interactions within the NIC phenyl rings of NIC [35,36].

However, MgO being basic in nature, could induce deprotonation of phenolic proton in NIC. The two well-known monohydrates such as the alpha and beta conformations of NIC are associated with the phenolic proton [37]. The alpha conformation associated with hydrogen bonding between phenolic proton and carbonyl group, while the beta conformation associated with phenolic and amido proton (Figure 5).

**Fig. 5.**
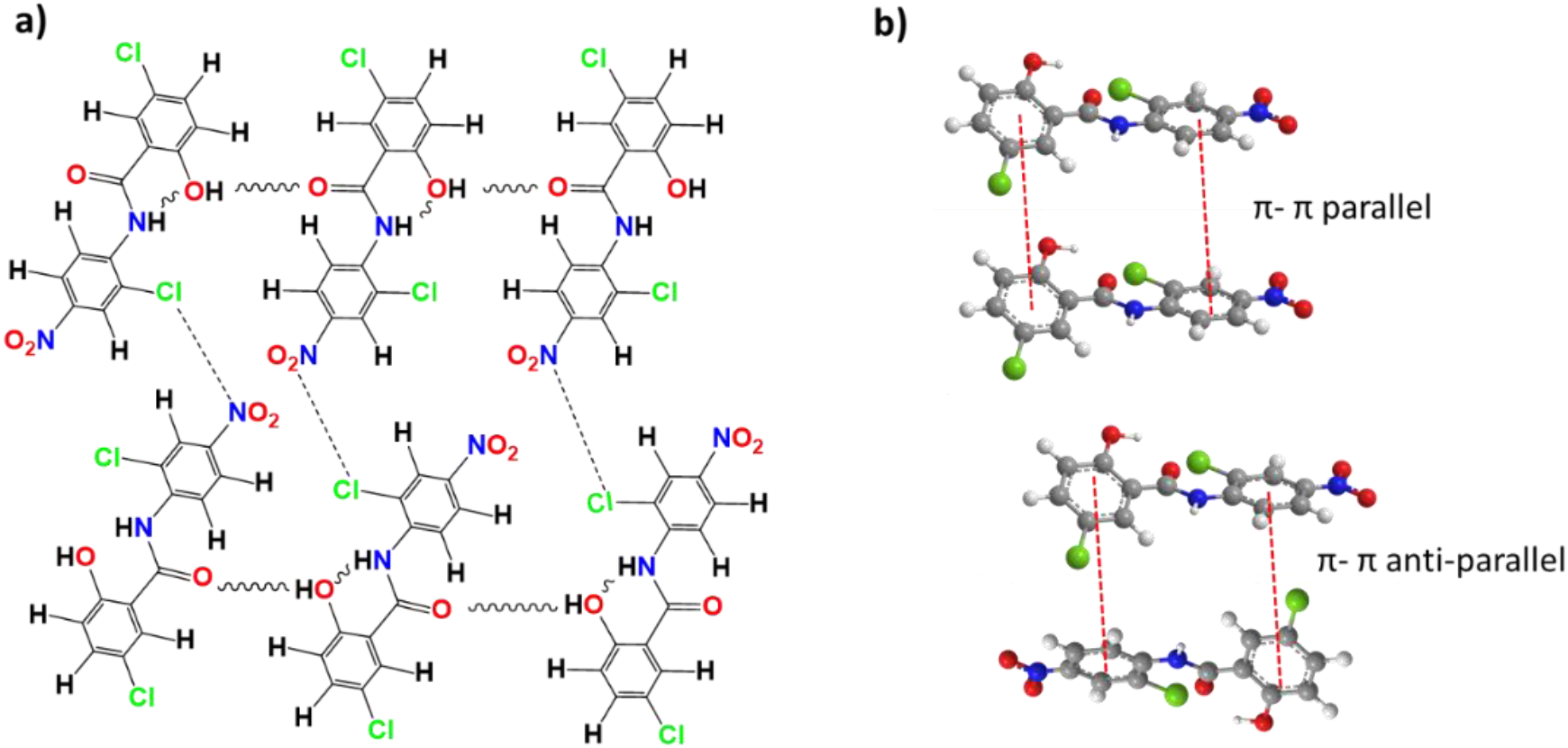
Molecular interactions in single crystals of NIC a) inter and intra molecular hydrogen bonding; b) π-π parallel and anti-parallel interactions between the NIC layers

This deprotonation therefore induce an anion-cation anti-parallel interaction thereby breaking the π-π stacks between the NIC molecules in the NIC-MgO structure, making them more water soluble, as previously suggested by Vuai et al [35]. Similar cation-anion interaction between NIC molecules were previously reported [36]. This structure model was further evidenced by UV analysis (Figure 6). The solid UV analysis indicated the specific bands at 368, 352 and 285 nm for the NIC, of which both 368 and 352 bands were shifted to 352 and 310 after a hydrogen bonding interaction with MgO. The 285 band was further shifted to 237 nm, clearly indicating the involvement of strong hydrogen bonding. In ethanol solution, NIC molecules were distinguished by absorption bands at 211 and 333 nm. In the solid NIC, the packing of the NIC molecules were very close and regular and the interactions of aromatic groups could result in π-π stacking, which leads to a red shift of the absorption bands at 285 nm and 352 nm. Furthermore, a blue shift to 237 nm and 310 nm in the NIC-MgO could be due to the breaking of π-π stacks between the NIC molecules and increasing a strong interaction between NIC and MgO in their composite form. Identical absorption bands were also obtained from the as-made NIC-MgO-HPMC, which demonstrated the formation of broken π-π stacking interactions [38].

**Fig. 6.**
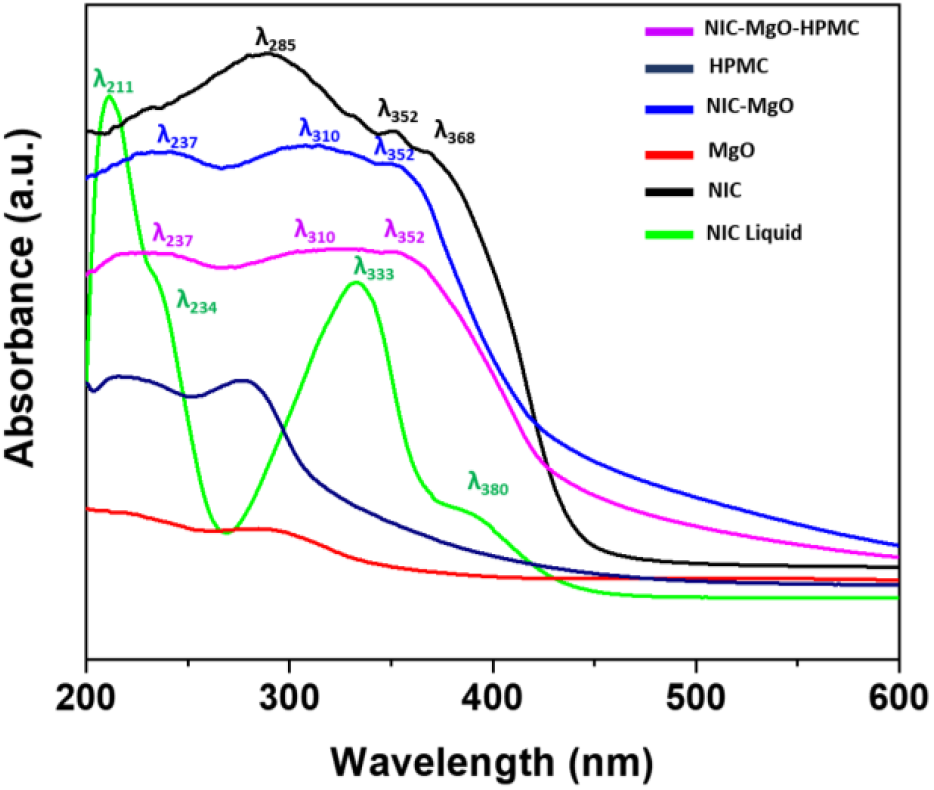
Solid UV-Vis absorption spectra of NIC, NIC-MgO, HPMC, NIC-MgO-HPMC and liquid UV-Vis absorption spectrum of NIC in Ethanol solution

The greatest challenge in NIC delivery is its low aqueous solubility as previously reported by many researchers, including our group by repurposing technology using polymers [39, 40] as well as inorganic clays [41].

The solubility of NIC is partly dependent on the self-association in water. To clearly understand this issue of aggregated NIC, four different interaction modes have been considered in classical MD simulations of different cluster sizes. It was found that the low water solubility of NIC is due to the antiparallel arrangements characterized by the formation of the cation–π and π–π interactions. Therefore, the solubility of NIC could be increased only by breaking the cation–π and π–π interaction by introducing stronger cation-anion interactions [36].

Prior to the release, the drug contents were measured, where the NIC-MgO had ∼ 56.3 ± 3.4 % drug content while the NIC-MgO-HPMC had ∼ 49.6 ± 1.4 % respectively (Table 2). As expected, the NIC-MgO-HPMC improved the solubility which resulted in to enhanced drug release, especially in pH 6.8 than that of pH 1.2, and the release was much higher than the counterparts as shown in Figure 3. In pH 6.8, the intact NIC showed a dissolution of 55%, whereas the commercial NIC formulation, Yomesan, had ∼ 63% NIC release, both of them showed low drug release (**Fig. 7**)

**Table 2.**
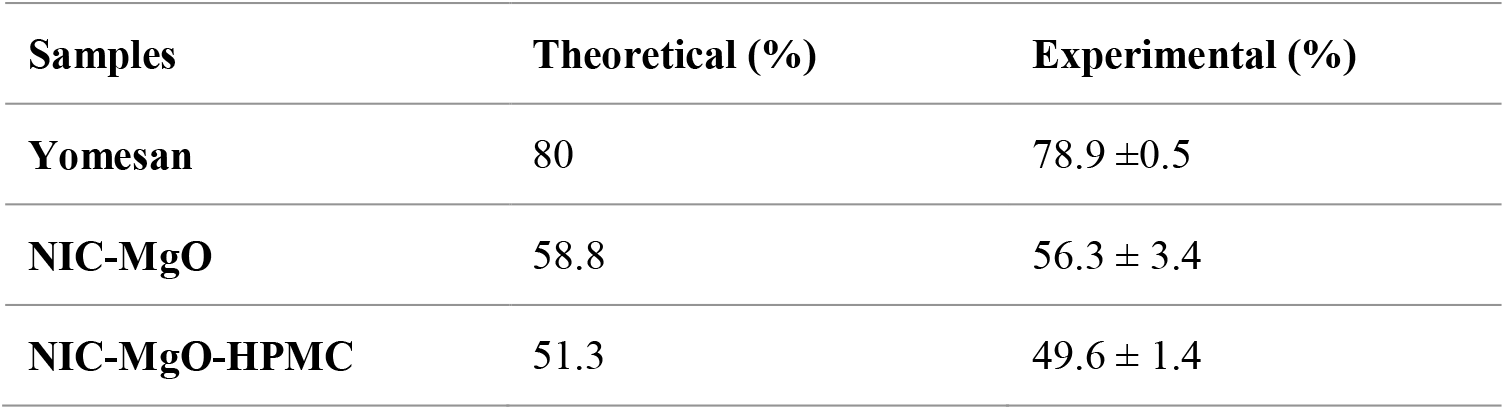
The NIC content of Yomesan, NIC-MgO, NIC-MgO-HPMC

| Samples | Theoretical (%) | Experimental (%) |
| --- | --- | --- |
| Yomesan | 80 | 78.9 ± 0.5 |
| NIC-MgO | 58.8 | 56.3 ± 3.4 |
| NIC-MgO-HPMC | 51.3 | 49.6 ± 1.4 |

**Fig. 7.**
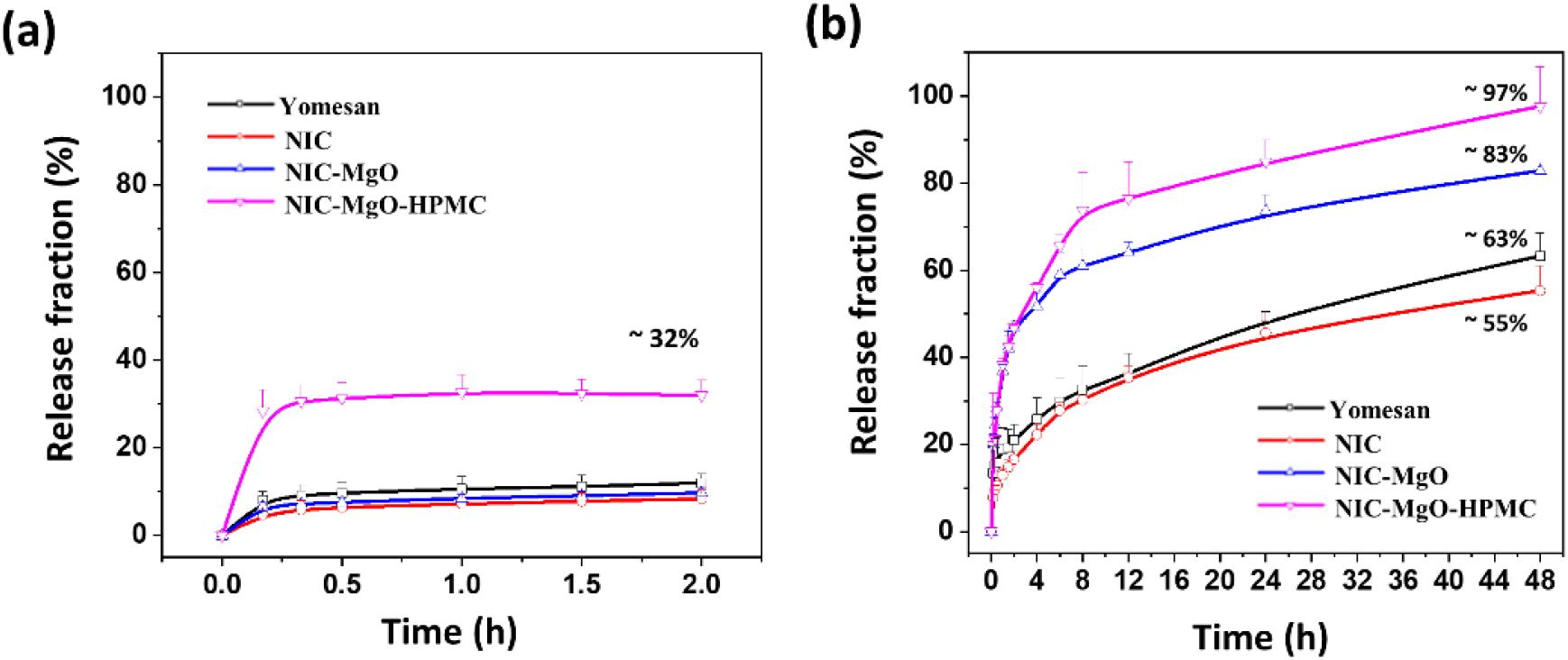
*In vitro* release test of Yomesan, intact NIC, NIC-MgO and NIC-MgO-HPMC in (a) pH 1.2 medium and (b) pH 6.8 medium that contained 2% Tween 80

Generally, the stable conformation for the NIC crystals are stabilized by the formation of cation– π stacking interactions that, on increasing the cluster size, the inter planar distances of parallel molecules lowered, indicating the possible reason behind the poor solubility of NIC. It has been reported that, the β-conformation of NIC exists in its planar conformation stabilized through an intramolecular hydrogen bond between the amido and hydroxyl protons [36]. It is therefore expected that both NIC and Yomesan could prefers to be in antiparallel conformation, while the NIC-MgO and NIC-MgO-HPMC could be in parallel conformation as well [36].

The improved NIC drug release from NIC-MgO-HPMC could be due to the combined effects of MgO and HPMC coating. However, in all the cases, there was restricted NIC release found at acidic pH (< 40%) (Fig. 7). This low release could be due to the selectivity of HPMC as it is more prone to pH > 3 ∼ 3.4, than that at pH 1.2 [42]. In addition, the initial burst release at GIT could be heavily limited by the HPMC coating, while the acid labile MgO can further sustain the release in the intestinal pH condition. This is because MgO could undergo ∼ 80% dissolution in acidic pH just within 30 min, while at neutral pH, MgO is could be completely soluble in slower pace of 48h (Fig. S1), which is very important for sustaining NIC release at the intestinal pH conditions. Further, we have studied the time dependent pH variations (Fig. S2), and was found that only MgO could increase the pH to nearly basic, thereby improving the drug solubility. It has been reported that NIC solubility could be improved with increasing pH from 7 to 9.3 [43]. Further experiments on pH variations of samples over different time periods were studied at pH 1.2 and 6.8, clearly showing that presence of MgO could further enhance the –OH concentration under a pH of 6.8, allowing improved solubility.

The amounts of remaining NIC after reaction with intestinal microsome at various time intervals were plotted in Fig. 8. With rat intestinal microsome, NIC levels were decreased over time upon incubation with either NADPH or UDPGA, while a negligible metabolism occurred in the absence of UDPGA. Co-incubation with both cofactors (NADPH and UDPGA) led to a further decrease in remaining NIC amount. With intestinal microsome, more than 80% of NIC disappeared within 15 min in the presence of NADPH and UDPGA. The rates of decrease of NIC were similar for NIC and NIC-MgO-HPMC suggesting that the NIC formulation with MgO and HPMC does not affect the metabolism of NIC.

**Fig. 8.**
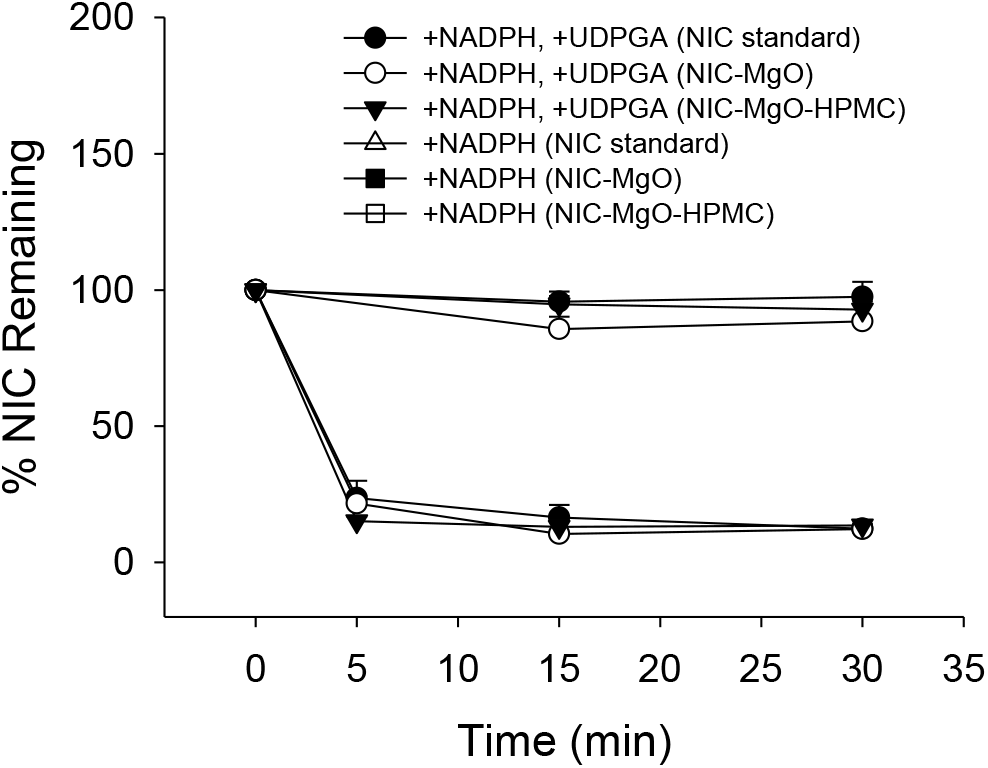
*In vitro* metabolic stability of NIC standard, NIC-MgO, and NIC-MgO-HPMC in rat intestinal microsomes with and without UDPGA treatment.

PK analyses were performed on hamster model using various concentrations of NIC-MgO-HPMC such as 20, 40 and 80 mg/kg and was compared with NIC (80 mg/kg) to understand the difference between commercially available NIC product with the repurposed one (Fig. 9a). As expected, the PK performance was dependent on the dosage as we could observe an improved AUC as the dosage increased from 20 to 80 mg/kg (Fig.9b). In addition, the PK parameters such as C max, t _1/2_, t max and AUC were improved with the as made repurposed formulation (Fig. 9c), clearly indicating their potential efficacy as anti-COVID-19 drug candidate.

**Fig. 9.**
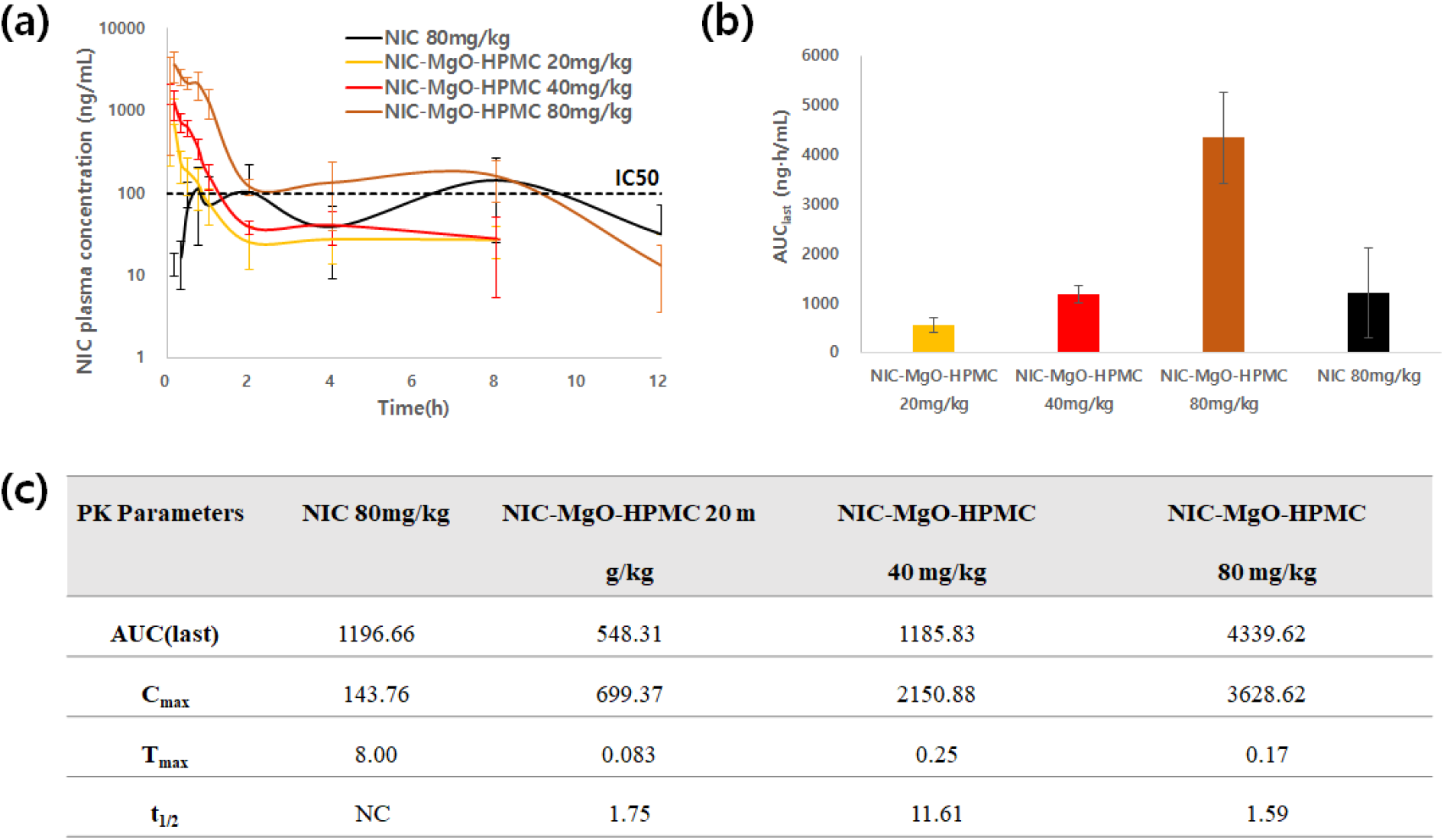
PK profile of NIC by NIC-MgO-HPMC formulation. (a) *In vivo* PK profiles of NIC and NIC-MgO-HPMC and (b) AUC derived from PK profiles of NIC and NIC-MgO-HPMC (n=5, ** represents p<0.05). (c) PK parameters of NIC (80 mg/kg), and NIC-MgO-HPMC at 20, 40 and 80 mg/kg doses in golden Syrian hamster model (n= 5) (mismatching table and PK Data in t_1/2_)

Most importantly, the NIC-MgO-HPMC at higher dosage of 80 mg/kg was able to sustain the NIC plasma concentration even up to 8 h constantly in a sustained manner from 2 h till 8h, which was above the required IC50 value for NIC. The lower dosages were however declined to below the IC50 values after 2h. This indicates that higher dosages of NIC-MgO-HPMC would be a good option than the lower ones.

As shown in Fig.10a, NIC-MgO-HPMC was orally administered to SARS-Co2-infected golden syrian hamster. At 4 dpi, all animals were sacrificed to collect the lung tissues for quantitative analysis on viral RNA levels by RT-qPCR. Compared to vehicle-treated hamsters (VC), NIC-MgO-HPMC treatment lowered rate of SARS-CoV2 replications in the lung tissues. Even though lung viral loads were reduced in individual lung lobes after NIC-MgO-HPMC treatment, this was found to be statistically relevant only in few lung lobes, owing to higher variation between the animals in the therapeutically treated group (Fig.10b). The necropsy evaluation on each area of lung lobe infected by SARS-CoV-2 was quantified through a specific score assignment. Gross lung lesions were observed in several lung lobes of vehicle-treated control group (VC). On the other hand, the animals treated with 40 or 80mg of NIC-MgO-HPMC exhibited lower gross lung lesions (Fig. 10c). Additionally, the total area of lungs with gross lesions was significantly lowered than in VC (Fig.10d). Further, the analysis on severity of histologic lung lesions was also studied and the resulting lung histology score was compared between treatment groups to understand differences in the severity of histologic lesions. The 80mg NIC-MgO-HPMC treated animals exhibited lower lung histology scores indicating the safety of the oral formulation (Fig. 10e). Further, toxicity of orally administered NIC-MgO-HPMC was studied in both rodents and non-rodents. In the repeated-dose toxicity study in which NIC-MgO-HPMC was administered to rats, no mortality or abnormal symptoms were observed. In body weight, feed intake, ophthalmological examination, urinalysis, haematological examination, organ weight, autopsy and histopathological examinations, significant changes were not observed. Therefore, the NOAEL (No-observed-adverse-effect level) of NIC-MgO-HPMC is considered to exceed 800 mg/kg in both sexes in rats (Table 3).

**Table 3.**
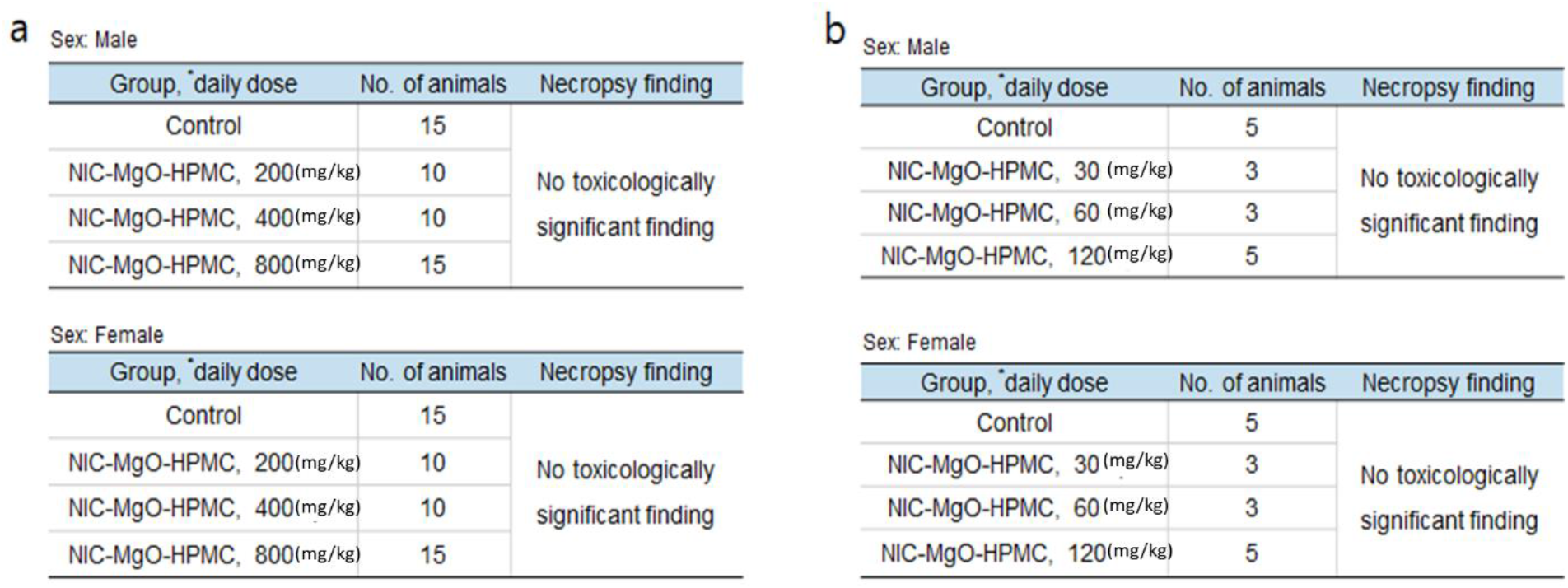
Toxicity study results. Repeated dose toxicity using (a) rats and (b) beagles.

**Fig. 10.**
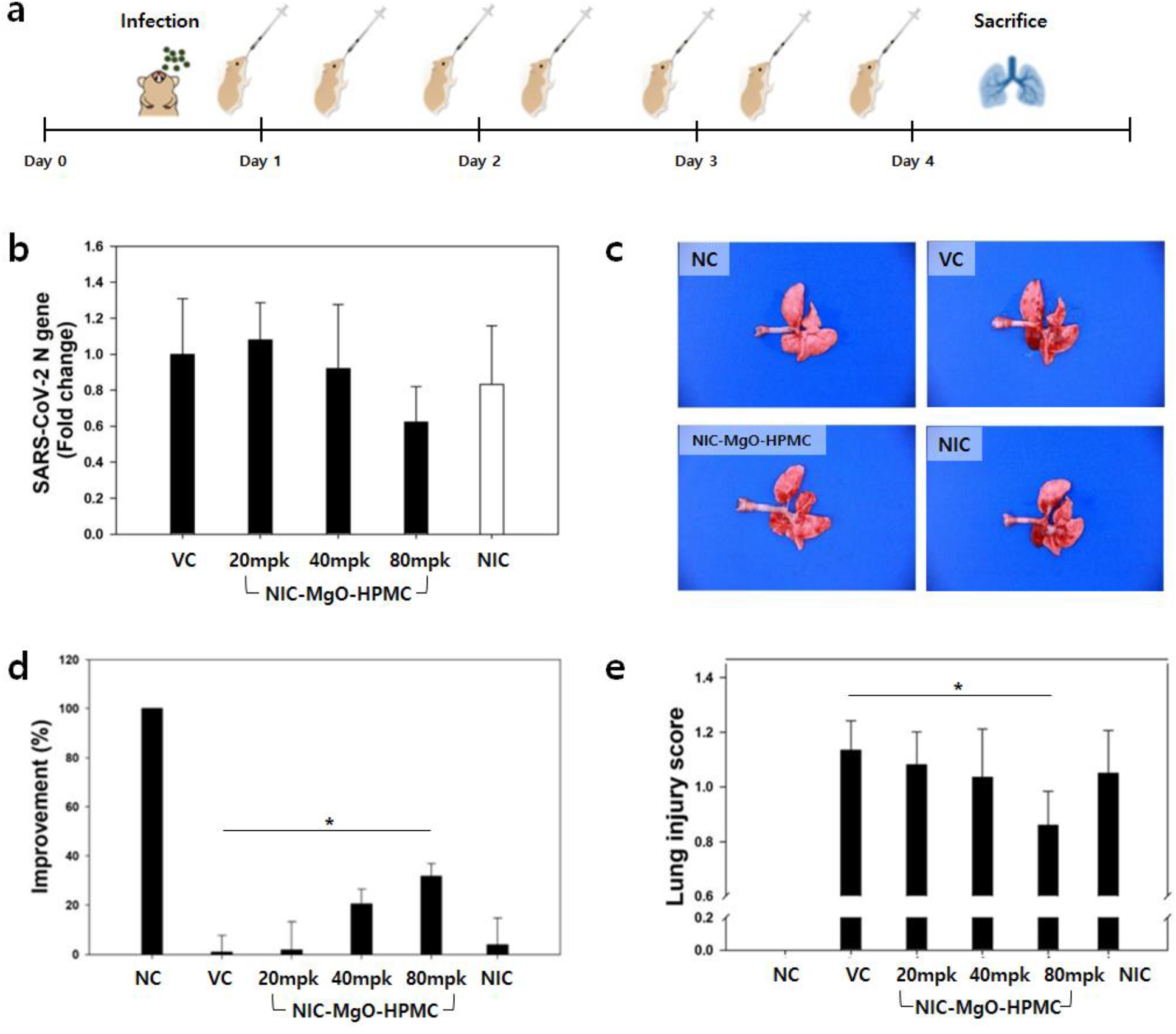
Therapeutic activity of NIC-MgO-HPMC observed in golden Syrian hamster model. (a) Hamsters were intranasally infected with SARS-CoV-2. NIC-MgO-HPMC was administered post-infection starting 6h post-infection. Treatment was continued every 12h for 4 days. Hamsters were euthanized on 4 dpi and lungs were harvested. (b) Lung viral loads were determined by using RT-qPCR. (c) One representative image of lung was chosen for each group. (d) Gross pathological lung lesion. At necropsy, the percentage of each lung lobe affected by gross lesions was estimated. (e) Histopathological examination of lungs of hamster model. Lung samples were stained with H&E and analyzed for the presence of lesion.

In a toxicity test conducted by repeated administration of NIC-MgO-HPMC to beagle, no deaths were observed in both males and females. During the administration period, symptoms such as vomiting and diarrhoea were observed in a dose-dependent manner in all dose groups of both sexes. However, these symptoms were immediately recovered when the administration of NIC-MgO-HPMC was stopped. In addition, as no toxicity was observed in body weight, feed intake, eye examination, electrocardiogram organ weight, and histopathological examination, the NOAEL for male and female beagle dogs is considered to exceed 120mg/kg. To calculate the maximum recommended starting dose (MRSD) in entry into human studies, the NOAEL values in animal species can be converted into human equivalent dose (HED). We calculated MRSD using NOAEL in both species by applying allometric scale of rat (6.2) and beagle (1.8) (Table 3).

## 4. Discussion

The current situation of pandemic is devastating as there has been not much significant decline in the overall daily cases of COVID-19. In addition, the surging new cases are associated with the rise of new variants such as the one reported in Indian subcontinent, the delta variant, causing millions affected, with lakhs dead [44-47]. Scientists across the world are more concerned about these new variants [48, 49]. Therefore, it is an urgent need to develop new medications which could be combined with the ongoing global vaccination strategies to have better effects. Additionally, the G7 summit held in Germany on May 31^st^ 2022, urged for an advance preparation of broad spectrum antiviral for facing the next pandemic in the near future. Accordingly, NIH and other major company associations have invested huge amount of 577 million US$ to initiate the research outputs up to phase I clinical trials.

In this context, it is an urgent need for a master drug formulation which could be useful for wide range of viral species. Here, our repurposed NIC formulation has all the ingredients as FDA approved ones, including the MgO and HPMC, making it as safe oral drug formulation towards COVID-19. The analysis results revealed their possible interaction within the composite majorly *via* hydrogen bonding and are able to have a predominant release at the intestinal pH compared to that at gastric one. This is very crucial as the SARS-CoV-2 virus can easily infect not only the respiratory cells [50, 51], but also intestinal organs [52-54], and would be highly effective in treating those who are in the late state of viral infection. The better release performance of NIC from the composite would be explained by the preferential dissolution of HPMC at pH > 3, while the MgO could undergo much slower dissolution at the same pH condition, thereby the encapsulated NIC could have a sustained release over 48 h. Here, the idea of HPMC coating is to restrict the burst release during the GIT entry and also to inhibit the harsh acidic conditions, so that NIC would only be released at the specific targeted site, where one can localize more NIC to have better efficacy. Thus, the rationally designed NIC-MgO-HPMC could be highly beneficial. Further PK analysis revealed that NIC can be sustained for longer duration at the dosage of 80 mg/kg, and was much higher than that of the control NIC formulation that is commercially available. PK analysis of NIC in hamster model displayed a linear increase in plasma exposure (AUC) between the doses.

Further *in-vivo* anti-SARS-CoV2 analysis on hamster model confirmed that the as made oral pill could be effective in containing the virus as it was effective than that of NIC itself. In our current study, the well-established Syrian hamster model was utilized to confirm the *in vivo* anti-viral efficacy of NIC-MgO-HPMC on SARS-CoV-2 replication. NIC-MgO-HPMC reduced the replication of SARS-CoV-2 in the lungs based on both viral RNA genome copy number determined by RT-qPCR. Importantly, the reduced replication of the viruses was associated with markedly reduced lung pathology. The high dose of NIC-MgO-HPMC used in this study (80 mg/kg BID) resulted in the 40% reduction of viral load in lungs, improvement in gross pathological lesion and histopathological scores in lungs.

How clinically realistic are these results? To address this question, we need to convert the hamster dose to HED. The therapeutic dose (80 mg/kg) of NIC-MgO-HPMC obtained in the hamster model is corresponding to 10.8 mg/kg in HED. Considering that the approved maximum daily dose of NIC is 2g (33.3mg/kg), it is estimated that NIC-MgO-HPMC can have a therapeutic effect within a realistic dose. Additionally, the NOAEL (> 800 mg/kg in rat, >120 mg/kg in beagle) observed in rodent and non-rodent animals support that the therapeutic dose is lower than the toxic dose. The safety and PK of NIC-MgO-HPMC were investigated in healthy adult participants and now, a clinical trial phase 2 is currently underway in Korea under the drug name CP-COV03. The results of the clinical trial will be additionally reported in the future.

## Acknowledgements

This research was supported by Basic Science Research Program through the National Research Foundation of Korea (NRF) funded by the Ministry of Education (No. 2020R1I1A2074844), and under the framework of the International Cooperation Program managed by NRF (2017K2A9A2A10013104).

## Appendix

A. Supplementary data

The following are the supporting information attached:

**Fig. S1** Solubility of MgO in (a) pH 1.2 and (b) pH 6.8 media. (Accordingly the figure numbers will be changed hereafter)

**Fig. S2** Time dependent pH variations of NIC, NIC-MgO, and NIC-MgO-HPMC under release conditions using 2% Tween 80 continuing release media) pH 1.2 and b) 6.8.

## References

[1] T.V. Padma. How COVID changed schools outreach. Nature 594(2021) 289–291. 10.1038/d41586-021-01517-7

[2] S. Mallapaty. The COVID vaccine pioneer behind southeast Asia’s first mRNA shot. Nature 594(2021)163. DOI: https://doi.org/10.1038/d41586-021-0142

[3] S-P, Jun, H.S. Yoo, J-S. Lee. The impact of the pandemic declaration on public awareness and behavior: Focusing on COVID-19 google searches. Technological Forecasting and Social Change 166(2021)120592. https://doi.org/10.1016/j.techfore.2021.120592

[4] P. Mlcochova, S.A. Kemp, M.S. Dhar, et al. SARS-CoV-2 B.1.617.2 Delta variant replication and immune evasion. Nature 599(2021)114–119. https://doi.org/10.1038/s41586-021-03944-y

[5] J.R.C. Pulliam, C. V. Schalkwyk, N. Govender., et al. Increased risk of SARS-CoV-2 reinfection associated with emergence of the Omicron variant in South Africa. Science. 376 (2022)6593. DOI: 10.1126/science.abn494.

[6] F. Kreier. Deltacron: the story of the variant that wasn’t. Nature, 602(2022)19. DOI: https://doi.org/10.1038/d41586-022-00149-9

[7] V. T. Vitiello, R. L. Porta. What will be the role of molnupiravir in the treatment of COVID-19 infection? Drugs Ther. Perspect. 37(2021) 579–580. https://doi.org/10.1007/s40267-021-00879-2

[8] Willyard. How antiviral pill molnupiravir shot ahead in the COVID drug hunt. Nature, (2021). DOI: https://doi.org/10.1038/d41586-021-02783-1

[9] E. Mahase. Covid-19: Molnupiravir reduces risk of hospital admission or death by 50% in patients at risk, MSD reports. BMJ 375 (2021) 2422. DOI: 10.1136/bmj.n2422.

[10] W. Fischer et al. Molnupiravir, an Oral Antiviral Treatment for COVID-19. medRxiv, (2021) DOI: 10.1101/2021.06.17.21258639

[11] B. Ahmad, M. Batool, Q.U. Ain, M.S. Kim, S. Choi. Exploring the Binding Mechanism of PF-07321332 SARS-CoV-2 Protease Inhibitor through Molecular Dynamics and Binding Free Energy Simulations. Int. J. Mol. Sci. 22 (2021) 9124. DOI: 10.3390/ijms22179124.

[12] M. Pavan, G. Bolcato, D. Bassani, M. Sturlese, S. Moro. Supervised molecular dynamics (SuMD) Insights into the mechanism of action of SARS-CoV-2 main protease inhibitor PF-07321332. J Enzyme Inhib Med Chem 36 (2021)1646–1650. DOI: 10.1080/14756366.2021.1954919

[13] S. Zhou, C.S. Hill, S. Sarkar., et al. β-d-N4-hydroxycytidine Inhibits SARS-CoV-2 through lethal mutagenesis but is also mutagenic to mammalian cells. J. Infect. Dis. 224 (2021) 415–419. DOI: 10.1093/infdis/jiab247

[14] R. Golubchikova, I. Danilycheva. Antihistamine therapy efficacy in patients with chronic idiopathic urticaria. Russ. J. Allergy. 9 (2012)13–18.

[15] N.C. Gassen, D. Niemeyer, D. Muth, et al. SKP2 attenuates autophagy through Beclin1-ubiquitination and its inhibition reduces MERS-Coronavirus infection. Nat. Commun. 10(2019) 5770. https://doi.org/10.1038/s41467-019-13659-4

[16] N.C. Gassen, J. Papies, T. Bajaj, et al. SARS-CoV-2-mediated dysregulation of metabolism and autophagy uncovers host-targeting antivirals. Nat. Commun. 12 (2021)3818. https://doi.org/10.1038/s41467-021-24007-w

[17] C-K . Lin, M-Y Bai, T-M Hu., et al. Preclinical evaluation of a nanoformulated antihelminthic, niclosamide, in ovarian cancer. Oncotarget 7(2016)8993–9006. DOI: 10.18632/oncotarget.7113

[18] R. Ma, Z-G. Ma, J-L. Gao., et al. Injectable pegylated niclosamide (polyethylene glycol-modified niclosamide) for cancer therapy. J. Biomed. Mater. Res. A 108(2020)30–38. DOI: 10.1002/jbm.a.36788

[19] S. Shah, M.M. Dooms, S. Amaral-Garcia, M. Igoillo-Esteve. Current drug repurposing strategies for rare neurodegenerative disorders. Front. Pharmacol. 12(2021)768023. DOI: 10.3389/fphar.2021.768023

[20] G. Choi, H. Piao, Z.A. Alothman, A. Vinu, C-O. Yun, J-H. Choy. Anionic clay as the drug delivery vehicle: tumor targeting function of layered double hydroxide-methotrexate nanohybrid in C33A orthotopic cervical cancer model. Int J Nanomed 11(2016)337–348. DOI https://doi.org/10.2147/IJN.S95611

[21] J-M. Oh, M. Park, S-T. Kim, J-Y. Jung, Y-G. Kang, J-H. Choy. Efficient delivery of anticancer drug MTX through MTX-LDH nanohybrid system. Journal of Physics and Chemistry of Solids 67(2006)1024–1027. https://doi.org/10.1016/j.jpcs.2006.01.033

[22] G. Choi, et al. Hydrotalcite–niclosamide nanohybrid as oral formulation towards SARS-CoV-2 viral infections. Pharmaceuticals 14 (2021) 486. https://doi.org/10.3390/ph14050486

[23] C.F. Tang, H. Ding, R.Q. Jiao, X.X. Wu, L.D. Kong. Possibility of magnesium supplementation for supportive treatment in patients with COVID-19. Eur. J. Pharmacol. 886(2020) 173546 DOI: 10.1016/j.ejphar.2020.173546

[24] B. Mishra, J. Sahoo, P.K. Dixit. Formulation and process optimization of naproxen nanosuspensions stabilized by hydroxy propyl methyl cellulose. Carbohydr Polym 127, (2015)300–308. DOI: 10.1016/j.carbpol.2015.03.077

[25] J. Vijay, J. Sahadevan, R. Prabhakaran, R.M. Gilhotra. Formulation and evaluation of cephalexin extended-release natrix tablets using hydroxy propyl methyl cellulose as rate-controlling polymer. J .Young. Pharm. 4(2012)3–12. DOI: 10.4103/0975-1483.93570

[26] V.V. Thai, B.T. Lee. Fabrication of calcium phosphate-calcium sulfate injectable bone substitute using hydroxy-propyl-methyl-cellulose and citric acid. J Mater Sci Mater Med 21 (2010)1867–1874. DOI: 10.1007/s10856-010-4058-9

[27] Arezzo, et al. Hydroxy-propyl-methyl-cellulose is a safe and effective lifting agent for endoscopic mucosal resection of large colorectal polyps. Surg Endosc 23 (2009)1065–1069. DOI: 10.1007/s00464-008-0133-4.

[28] A. Foppoli, A. Maroni, L. Palugan, et al., Erodible coatings based on HPMC and cellulase for oral time-controlled release of drugs. Int J Pharm. 585(2020)119425. doi: 10.1016/j.ijpharm.2020.119425.

[29] S. Yu, H. Piao, N.S. Rejinold, G-W. Jin, G. Choi, J-H. Choy. Niclosamide–clay intercalate coated with nonionic polymer for enhanced bioavailability toward COVID-19 treatment. Polymers 13 (2021) 1044. https://doi.org/10.3390/polym13071044

[30] H. Piao, N.S. Rejinold, G. Choi, Y-R. Pei, G-W. Jin, J-H. Choy. Niclosamide encapsulated in mesoporous silica and geopolymer: A potential oral formulation for COVID-19. Micropor Mesopor Mat 326(2021)111394 . https://doi.org/10.1016/j.micromeso.2021.111394

[31] K. Rosenke, et al. Orally delivered MK-4482 inhibits SARS-CoV-2 replication in the Syrian hamster model. Nat. Commun. 12(2021)2295. https://doi.org/10.1038/s41467-021-22580-8

[32] D.R. Owen, et al. An oral SARS-CoV-2 M pro inhibitor clinical candidate for the treatment of COVID-19. Science, 374 (2021) 1586–1593. DOI: 10.1126/science.abl478

[33] Ahmad, et al. Self-assembly and wetting properties of gold nanorod-CTAB molecules on HOPG. Beilstein journal of nanotechnology 10(2019)696–705

[34] D. Maity, et al. Synthesis of HPMC stabilized nickel nanoparticles and investigation of their magnetic and catalytic properties. Carbohydr. Polym. 98(2013)80–88. DOI: 10.1016/j.carbpol.2013.05.020

[35] S.A.H. Vuai, M.G. Sahini, I. Onoka, L.W. Kiruri, D.M. Shadrack. Cation–π interactions drive hydrophobic self-assembly and aggregation of niclosamide in water. RSC Adv. 11(2021)33136–33147. https://doi.org/10.1039/D1RA05358B

[36] P. Sanphui, S.S. Kumar, A. Nangia. Pharmaceutical cocrystals of niclosamide. Crystal Growth & Design. 12 (2012)4588–4599. https://doi.org/10.1021/cg300784v

[37] D. Luedeker, R. Gossmann, K. Langer, G. Brunklaus. Crystal engineering of pharmaceutical co-crystals: “NMR crystallography” of niclosamide co-crystals. Crystal Growth & Design 16 (2016) 3087-3100. doi/10.1021/acs.cgd.5b01619

[38] D. Xu, et al. π–π interaction of aromatic groups in amphiphilic molecules directing for single-crystalline mesostructured zeolite nanosheets. Nat. Commun. 5 (2014)4262. https://doi.org/10.1038/ncomms5262

[39] N.S. Rejinold, H. Piao, G-W. Jin, G. Choi, J-H. Choy. Injectable niclosamide nanohybrid as an anti-SARS-CoV-2 strategy. Colloids Surf B Biointerfaces 208(2021)112063. DOI: 10.1016/j.colsurfb.2021.112063

[40] N. S. Rejinold, G. Choi, H. Piao, J-H. Choy. Bovine serum albumin-coated niclosamide-zein nanoparticles as potential injectable medicine against COVID-19. Materials 14, (2021)3792. DOI: 10.3390/ma14143792

[41] N.S. Rejinold, H. Piao, G. Choi, G-W. Jin, J-H. Choy. Niclosamide-exfoliated anionic clay nanohybrid repurposed as an antiviral drug for tackling covid-19; oral formulation with tween 60/eudragit s100. Clays. Clay Mineral. 69(2021)533–546. DOI: 10.1007/s42860-021-00153-6

[42] Y. Huang, L.L. Zheng, J. Liu, Z.R. Zhang. Synthesis and characterization of HPMC derivatives as novel duodenum-specific coating agents. Archives of pharmacal research 28(2005) 364–369. DOI: 10.1007/BF02977806

[43] D. Needham. The pH Dependence of niclosamide solubility, dissolution, and morphology motivates potentially universal mucin-penetrating nasal and throat sprays for COVID19, its contagious variants, and other respiratory viral infections. bioRxiv, 2021.2008.2016.456531 (2021). doi: https://doi.org/10.1101/2021.08.16.456531

[44] F. Graham. Daily briefing: Why the delta variant spreads so fast. Nature. (2021). doi: https://doi.org/10.1038/d41586-021-02032-5

[45] Reardon S. How the Delta variant achieves its ultrafast spread. Nature. (2021). DOI: https://doi.org/10.1038/d41586-021-01986-w

[46] D. Planas, D. Veyer, A. Baidaliuk, et al. Reduced sensitivity of SARS-CoV-2 variant Delta to antibody neutralization. Nature 596(2021)276–280. https://doi.org/10.1038/s41586-021-03777-9

[47] E. Callaway. Delta coronavirus variant: scientists brace for impact. Nature 595 (2021) 17–18. DOI: https://doi.org/10.1038/d41586-021-01696-3.

[48] F. Graham Daily briefing: Omicron coronavirus variant puts scientists on alert. Nature, (2021) DOI: 10.1038/d41586-021-03564-6

[49] E. Callaway. Heavily mutated omicron variant puts scientists on alert. Nature. 600 (2021)21. DOI: https://doi.org/10.1038/d41586-021-03552-w

[50] L.E. Wee, J.Y. Tan, S.J. Chung, E.P. Conceicao, B. Hock Tan, I. Venkatachalam. Zero health-care-associated respiratory viral infections among solid organ transplant recipients: Infection prevention outcomes during COVID-19 pandemic. Am J Transplant 21(2021)2311–2313. DOI: 10.1111/ajt.16499

[51] B. Baby, A.R. Devan, B. Nair, L.R. Nath. The impetus of COVID -19 in multiple organ affliction apart from respiratory infection: Pathogenesis, diagnostic measures and current treatment strategy. Infect. Disord. Drug. Targets. 21(2021)514–526. 10.2174/1871526520999200905115050

[52] M. Hirayama, H. Nishiwaki, T. Hamaguchi, et al. Intestinal Collinsella may mitigate infection and exacerbation of COVID-19 by producing ursodeoxycholate. PLoS One 16(2021)e0260451. 10.1371/journal.pone.0260451

[53] M.J. McAllister, K. Kirkwood, S.C. Chuah, et al. Intestinal protein characterization of SARS-CoV-2 entry molecules ACE2 and TMPRSS2 in inflammatory bowel disease (IBD) and fatal COVID-19 infection. Inflammation, 45(2021)567–572. DOI: 10.1007/s10753-021-01567-z

[54] X. Wang, Y. Zhou, N. Jiang, Q. Zhou, W.L. Ma. Persistence of intestinal SARS-CoV-2 infection in patients with COVID-19 leads to re-admission after pneumonia resolved. Int J. Infect. Dis. 95 (2020)433–435. DOI: 10.1016/j.ijid.2020.04.063

