## supplementary file for "Orally administered niclosamide-based organic/inorganic hybrid suppresses SARS-CoV-2 infection"

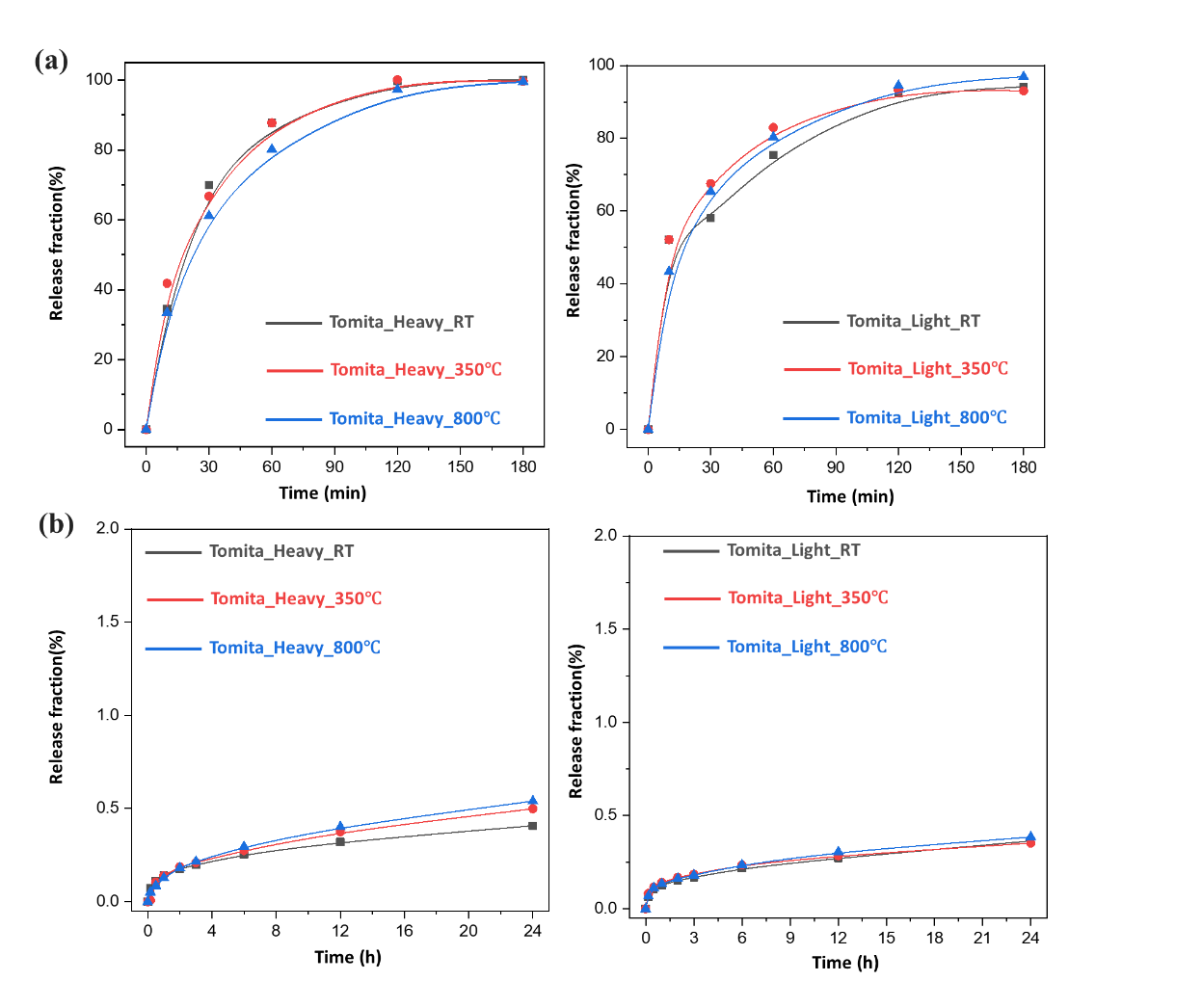

**Fig. S1** Solubility of MgO in (a) pH 1.2 and (b) pH 6.8 media. (Accordingly the figure numbers will be changed hereafter)

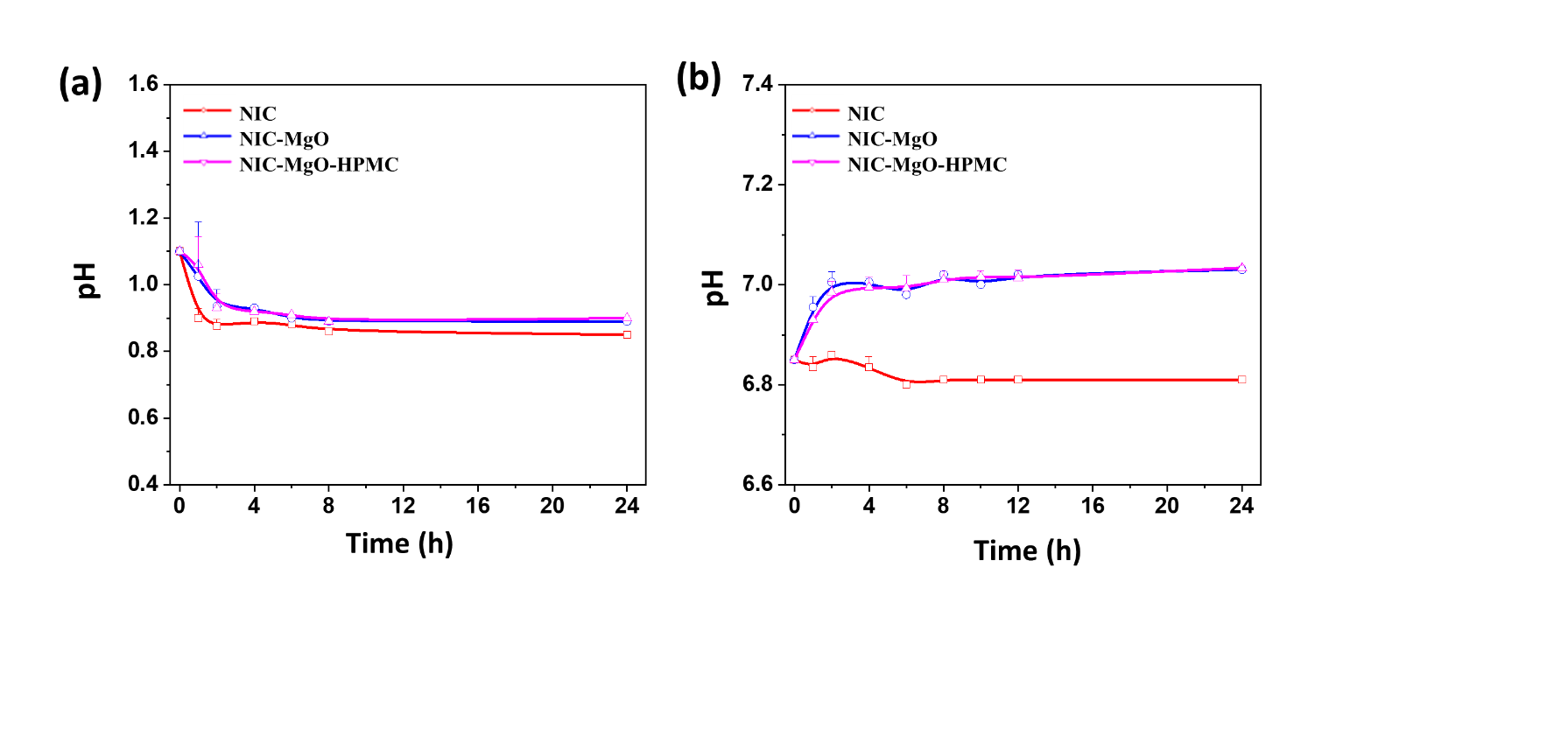

**Fig. S2** Time dependent pH variations of NIC, NIC-MgO, and NIC-MgO-HPMC under release conditions using 2% Tween 80 continuing release media) pH 1.2 and b) 6.8.
